# Lower neural value signaling in the prefrontal cortex is related to childhood family income and depressive symptomatology during adolescence

**DOI:** 10.1101/2021.01.13.426547

**Authors:** Esther E. Palacios-Barrios, Jamie L. Hanson, Kelly R. Barry, Dustin Albert, Stuart F. White, Ann T. Skinner, Kenneth A. Dodge, Jennifer E. Lansford

## Abstract

Lower family income during childhood is related to increased rates of adolescent depression, though the specific mechanisms are poorly understood. Evidence suggests that individuals with depression demonstrate hypoactivation in brain regions involved in reward learning and decision-making processes (e.g., portions of the prefrontal cortex). Separately, lower family income has been associated with neural alterations in similar regions. We examined associations between family income, depression, and brain activity during a reward learning and decision-making fMRI task in a sample of adolescents (full n=94; usable n=78; mean age=15.4 years). We identified neural regions representing 1) expected value (EV), the learned subjective value of an object, and 2) prediction error, the difference between EV and the actual outcome received. Regions of interest related to reward learning were examined in connection to childhood family income and parent-reported adolescent depressive symptoms. As hypothesized, lower activity in the subgenual anterior cingulate (sACC) for EV in response to approach stimuli was associated with lower childhood family income, as well as greater symptoms of depression measured one-year after the neuroimaging session. These results are consistent with the hypothesis that lower early family income leads to disruptions in reward and decision-making brain circuitry, which leads to adolescent depression.

## 1. Introduction

Child poverty is a prevalent societal problem, with detrimental effects for mental health (Reiss, 2013; Yoshikawa et al., 2012). In particular, adolescents from lower income families are more likely to experience elevated depressive symptoms, earlier onset of depressive episodes, and a more severe course of major depression (McLaughlin et al., 2011; Najman et al., 2010). Given these widely replicated findings, it is critical to identify mechanisms in order to predict, prevent, and treat depressive symptomatology after exposure to poverty. Such progress will likely emerge by: a) considering “multiple levels of analysis”, specifically neurobiological alterations related to exposure to poverty; and b) more precise consideration of the “lived experiences of poverty” and distinctions between facets of adversity (e.g., exposures and experiences).

In regards to neurobiology, adolescence is marked by significant structural and functional brain reorganization and growth (Casey, 2015; Giedd et al., 2015; Luna et al., 2010). Given the protracted developmental trajectories of areas like the amygdala and the hippocampus, it is perhaps not surprising that poverty and the stressors associated with lower family income have been related to alterations in these and connected brain areas (Johnson et al., 2016; Palacios-Barrios and Hanson, 2019). Less explored, however, have been potential alterations in the structure and function of the corticostriatal circuit, which include the ventral striatum (VS) and different sub-regions of the PFC, such as ventromedial PFC (vmPFC). Given the critical role of this brain circuit in motivation, reward responsiveness, and learning (Becker et al., 2019; Berridge and Robinson, 2003) and research finding that this brain circuit is often aberrant in depressed individuals (Forbes and Dahl, 2012; Pizzagalli, 2014), it may be particularly important to probe the functioning of this circuit in relation to poverty exposure and increased risk for depression.

As one thinks about poverty, it is critical to understand that “lived experiences” of poverty encompass a host of stressors and environmental disadvantages that pose threats to normative development. For instance, common stressors associated with poverty include harsher family interactions (e.g. inconsistent parental support; greater parental hostility) and community violence (Evans, 2004). However, conceptualizing exposure to poverty as a risk factor often blurs the distinction between “exposures” and “experiences”. In brief, exposures capture the probability of something occurring and the context that development occurs in (see (McLaughlin et al., 2020) for additional, thoughtful discussion of this issue). Exposures are, however, not a direct measurement of what a child actually experiences. Many youths may be exposed to different adverse developmental context (e.g., poverty), but actually not experience harsher family environments, community violence, and other negative experiences likely to directly impact development.

Related to potential brain alterations, past research has found that various forms of childhood adversity relate to behavioral correlates and neurobiological functioning in the corticostriatal circuit (Gianaros et al., 2011; Gonzalez et al., 2016; Marshall et al., 2018). For example, maltreated individuals have shown an impaired ability to update behavioral responses to rewards (Guyer et al., 2006) and slower learning of reward associations (Hanson et al., 2017; Sheridan et al., 2018). Neurobiologically, maltreatment and high levels of stressful experiences in childhood are related to lower activity in the VS (Hanson et al., 2016, 2015a) and smaller volumes and lower activity in medial portions of PFC (Fan et al., 2020; Van Harmelen et al., 2010). Notably, multiple reports from our research group have found that corticostriatal responsivity to positive feedback and rewards, but not negative feedback and punishment, is altered after exposure to high levels of childhood stress and adversity (Hanson et al., 2018, 2016, 2015a). Connected to these ideas, multiple reports in non-human animals have found rodents living in impoverished environments demonstrate features of anhedonia and aberrant processing of rewarding stimuli, including reductions in sucrose preference and peer-play (Bolton et al., 2018; Molet et al., 2016).

While the processing of positive feedback and rewards will be an important continued focus, it will also be critical to think about richly decomposing and parsing apart different component processes of the corticostriatal circuit. Indeed, basic cognitive neuroscience research indicates that the corticostriatal circuit underlies two important reward-learning and decision-making processes: the coding of prediction error (PE) and the estimation of value (or the expected value, EV). PE is the difference between an actual outcome and one’s expectations; EV is the “net value” associated with a stimulus and combines the magnitude of a reward with the potential probability of receiving it (Knutson et al., 2005; Rescorla and Wagner, 1972). These two aspects of reward learning and decision-making consistently activate regions within the corticostriatal circuit, with the VS being a key brain area for both PE and EV being strongly connected to functional patterns in portions of PFC, including: the subgenual cingulate (sACC), perigenual cingulate (pACC), and ventromedial PFC (vmPFC) (Garrison et al., 2013; O’Doherty, 2011; Rangel et al., 2008). Limited work has examined these types of processes in youth exposed to poverty or adverse experiences; the only notable exception focused on maltreated youths, which found these participants demonstrated blunted activity in the VS and the vmPFC during a passive avoidance decision-making task, compared to non-maltreated participants (Gerin et al., 2017). Although maltreatment was associated with alterations of these specific elements of the corticostriatal circuit, it is unclear whether exposure to poverty and lower family income may similarly influence neurobiology related to reward learning and decision-making. It may be possible that youth living in poverty are unable to robustly recruit corticostriatal hubs (e.g., VS; portions of the PFC) to track expected value across time and adapt behavioral responses according to such feedback. Thinking collectively, understanding how EV and PE, specifically in relation, to positive feedback and rewarding stimuli may be critical in understanding connections between poverty, depression, and neurobiology.

Connected to the “exposure” and “experience” distinction, we believe it will be critical to, first, probe the impact of exposures like poverty and lower socioeconomic status on brain development; indeed, millions of children are currently living in poverty and understanding broad impacts of this exposure are critical to public-health and public-policy (Shonkoff, 2016). We, however, believe such work should then, when possible, be followed up with a focus on specific adverse experiences common to poverty and likely to influence neurodevelopment. In this regard, a prominent model to more deeply understand experiences of poverty argues that environmental facets of harshness and unpredictability may differentially impact behavioral and neural development (Belsky et al., 2012; Ellis et al., 2009; McLaughlin et al., 2020).

Experiences of harshness are extrinsic threats to morbidity and mortality, while unpredictability involves uncertain and unexpected variations in developmental experiences, particularly unpredictable, chaotic, and harsh events (Baram et al., 2012; Ellis et al., 2009). Chronic occurrence of these types of experiences may adversely impact the cognitive skills and associated neurocognitive systems underlying reward learning and decision making. For example, harshness may impact the hypothalamic– pituitary-adrenal axis and stress-responsivity systems (Del Giudice et al., 2011) and this may then alter reward-related, dopaminergic functioning through glucocorticoid regulation (Kinner et al., 2016). Unpredictability, in contrast, may connect to volatile environmental input that then causes aberrant synaptic strengthening and pruning within the reward circuit (see (Birnie et al., 2020) for additional review). Both of these types of experiences are common in impoverished contexts. As such, considering reward learning and decision-making component processes (e.g., PE; EV), the broad exposure of child poverty, and also specific potential experiences common to these environments (e.g., harshness; unpredictability) may be particularly informative and important to understand links between development, neurobiology, and psychopathology.

Motivated by these ideas, we used a rich, prospective, longitudinal project (the Parenting Across Cultures Study; PAC) to examine associations between childhood family income, adolescent brain functioning related to reward learning and decision-making processes, and depression. We used a well-validated passive avoidance learning task that allowed us to probe PE and EV through computational modeling and neuroimaging (White et al., 2013). Based on past work, we hypothesized that lower family income in childhood would be related to lower functional activity: for EV within portions of the prefrontal cortex, and for PE in the VS, both for rewarding stimuli. We believe this lower activity is a sign of poor tracking and updating of choice values based on environmental feedback. We also hypothesized that these corticostriatal functional differences would be related to greater depressive symptoms assessed after the neuroimaging session. Finally, given that exposure to poverty often encompasses specific adverse experiences, we also explored potential associations between neurobiology and dimensions of adversity (e.g., harshness; unpredictability). Grounding our work in models put forth by Ellis and colleagues (e.g., (Ellis et al., 2017), as well as Baram and coworkers (Baram et al., 2012; Birnie et al., 2020), we conducted exploratory analyses to evaluate whether experiences of harshness versus unpredictability were related to functional brain activity during reward learning and decision-making.

## 2. Materials and Methods

### Participants

A subsample of youths and their parents from the Parenting Across Cultures (PAC) study was recruited for the current project. The PAC study is a prospective, longitudinal study of parenting practices and child development (see http://parentingacrosscultures.org for more details). Participants were recruited through local elementary schools and the full cohort consisted of 311 socioeconomically diverse families (109 European American, 103 African American, and 99 Hispanic). Youth participants and their parents were interviewed approximately every year, beginning when the participants were approximately 8 years of age. This project uses data from the beginning of the study through when participants were 16 years old, on average.

For this project, the original families were contacted to participate in a neuroimaging study. Of those that were contacted, 92 families chose to participate. The participants were approximately 15 years of age at the time of this neuroimaging session. Of the 92, 5 participants were excluded due to issues with computational modeling (e.g., poor model convergence; See “*Neuroimaging Task*” Section for description of model-fitting) and 9 participants were excluded due to fMRI modeling issues (e.g., no behavioral responses during one condition, causing inappropriate fMRI modeling fitting); no participants were excluded due to motion (>25% censored frames). The final (usable) sample, therefore, consisted of 78 participants (see Table 1 for descriptive statistics). Here, we connected neuroimaging to four PAC study time-points (Study Waves 2, 3, 5, and 7; neuroimaging was collected between Waves 5 and 6).

**Table 1.** Descriptive Statistics.

| **Variable** | **Statistic** |
| --- | --- |
| Sex | Female: 44.9% (N = 35); Male: 55.1% (N = 43) |
| Race/Ethnicity | European American: 44.9% (N = 35); African American: 35.8% (N = 28); Hispanic: 23.1% (N = 18) |
| Participant age (Study Start) | 9.02 (+/- 0.531) years, Range: 7.83 - 10.1 |
| Participant age (At Scanning Session) | 15.2 (+/- 0.671) years, Range: 13.6 - 16.9 |
| Participant age (Follow-up) | 16.4 (+/- 0.615) years, Range: 15.06 - 17.63 |
| Family Income (Midpoint of Income Bracket, Averaged Across 2 Years) | \$61,753.21 (+/- \$40,580), Range: \$5,000-\$115,500 |
| Depressive Symptom T-Score (~1 Year After the Neuroimaging Scan) | 54.546 (+/- 7.269), Range: 50-76 |
| CHAOS (Wave 5) | 13.454 (+/- 3.62), Range: 6-23 |
| Community Violence (Wave 5) | 2.649 (+/-3.54), Range: 0-18 |

PAC study time-points occurred approximately every year, but this was occasionally >12 months due to data collection logistics. A timeline is included in our Supplemental Materials that overviews this project in relation to the larger PAC study (*Figure S1*).

### Measures

#### Childhood family income

Parents used the Family Information Form to report family income when participants were approximately 10 and 11 years of age. The measure assesses family income on a 10-point scale ranging from 1-“up to $5,000” to 10-“beyond $81,000.” Parents selected an answer in response to the statement “Indicate the gross annual income of your family.” In keeping with past reports (Hanson et al., 2011), the midpoint of an income bracket selection was then log-transformed at each wave, and averaged for the two waves. For example, up to $5,000 had a midpoint of $2,500 and then a log-transformed value of 3.398; for the highest income bracket, a midpoint of $115,500 was used (bracketing between $81,000 and $150,000) and yielded a log-transformed value of 5.062.

#### Depressive Symptoms

Parents completed the Child Behavior Checklist (CBCL) one year after the neuroimaging scan (when participants were approximately age 16). The CBCL is a widely used caregiver-report used to assess child behavioral and emotional problems (Achenbach et al., 1991). Questions are scored on a three-point Likert scale (0 = “not true,” 1 = “somewhat or sometimes true,” 2 = “very true or often true”). Total raw scores for the Withdrawn/Depressed subscale were then standardized (to “T-Scores”); these compare the raw score to what would be typical compared to responses for youths of the same gender and similar age. Previous studies have shown that the Withdrawn/Depressed subscale strongly correlates with other measures of depression (Gomez et al., 2014; Kweon et al., 2016). This construct was also measured at the same time as family income (Wave 2, when participants were approximately 10 years of age); this earlier measure of withdrawn symptoms was used as a covariate in relevant analyses to limit the impact of early psychopathology on our variables of interest.

#### Exploratory Variables Focused on Environmental Harshness and Unpredictability

When adolescents were 15 years of age on average, parents completed a measure of harshness, as well as a measure of unpredictability. Related to harshness, parents completed a 7-item questionnaire on the perceived safety and livability of their neighborhood (O’Neil et al., 2001). Questions such as “I feel scared in my neighborhood” and “My neighborhood is a nice place to live” (reverse-coded) were rated using a four-point Likert scale (0 = “almost never true,” 4 = “almost always true”). Connected to unpredictability, parents completed the Confusion, Hubbub, and Order Scale (Matheny et al., 1995) to assess confusion, chaos, and disorder in their homes. Questions such as “It’s a real zoo in our home” and “The atmosphere in our home is calm” (reverse-coded) were rated on a five-point Likert scale (1 = “definitely untrue,” 4 = “definitely true”). Each measure was summed and entered into exploratory regression models as independent variables.

#### Neuroimaging Task

Participants completed a well-validated passive avoidance learning task during a neuroimaging scan (White et al., 2013). During the task, participants were presented with four stimuli, each associated with winning or losing points (a large amount or a small amount). On each trial, participants were first presented with one of four stimuli for 1500 ms and decided whether they wanted to actively “approach” it (via button press) or passively “avoid” it (withhold a response). Following the decision, a randomly jittered fixation cross was presented (0-4000 ms). Participants were then provided feedback for 1500 ms. Two of the stimuli resulted in winning points 70% of the time (one stimulus: 50 points; one stimulus: 10 points). The other two stimuli resulted in losing points 70% of the time (one stimulus: 50 points; one stimulus: 10 points). The high win (or lose) stimulus was always the same and would win (or lose) more points 70% of the time. Participants could only win or lose points if the stimulus was actively approached (via a button press). If participants chose to passively avoid the stimuli, a fixation cross was then presented instead for 1500 ms. Following the feedback, another randomly jittered fixation cross was presented (0-4000). Each stimulus was presented 14 times, for a total of 56 trials.

Each participant’s behavioral task data was used to model expected value (EV) and prediction error (PE) for each trial based on the Rescorla-Wagner model of learning (O’Doherty et al., 2007; Rescorla and Wagner, 1972). For the first trial of each object, EV was initially set to 0 and then updated using the formula:

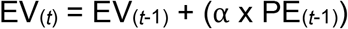

This formula indicates that the EV of the current trial (t) is equal to the EV of the previous trial (t-1) plus the PE of the previous trial multiplied by the learning rate. The learning rate was set to 0.667 after averaging across learning rates from all subjects that were estimated individually via a model-fitting simulation (see the Supplemental Methods section 47). The following formula was used to calculate the PE of the current trial:

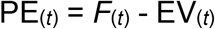

This formula indicates that the PE for the current trial is equal to the feedback (*F*) of the current trial minus the EV of the current trial. Parameters for EV and PE were extracted for both approached and avoided stimuli, and used a parametric regressors (“modulators”) in our model-based fMRI analyses (see *fMRI Data Preprocessing and Analysis* below).

### fMRI Data Acquisition

Scanning took place on a 3.0 Tesla General Electric scanner at the Duke-UNC Brain Imaging and Analysis Center. Structural and functional images were acquired during the scanning session. A high-resolution T1-weighted anatomical image was acquired, as well as whole-brain functional images were acquired using a SENSE inverse-spiral sequence (repetition time = 2000 ms; echo time = 32 ms; field of view = 256 mm; image matrix, 64×64; flip angle = 77 degrees; voxel size, 4×4×4 mm; 34 axial slices). Additional information about our neuroimaging acquisition are noted in our Supplemental Materials. Of note, resting state fMRI results from these same participants have been the focus of a previous publication (Hanson et al., 2019).

### fMRI Data Preprocessing and Analysis

Pre-processing and analysis of imaging data were conducted using Analysis of Functional Neuroimages (AFNI; http://afni.nimh.nih.gov, (Cox, 1996). Participants’ scans were realigned to the first acquisition volume. Individual fMRI time points were then censored if motion was >1.5 mm between frames, using the derivative and euclidean norm (through AFNI’s 1d_tool.py tool). We planned to exclude participants with ≥25% of frames censored; however, no participants met this threshold, meaning no participants were excluded for excessive motion. Data were then normalized into MNI-152 space using deformation fields from subjects’ T1-weighted scan, with a final voxel size of 2mm^3^. The resulting images were smoothed with a 6-mm Gaussian filter.

First-level individual analyses for each participant were calculated by convolving the event timing with the canonical hemodynamic response function modeling the four conditions: stimulus approached, stimulus avoided, reward received, punishment received. First-level models included parametric modulators (EV and PE) from the computational model, as well as nuisance covariates of the second-order polynomial used to model the baseline and slow signal drift, six motion estimate covariates and binary flags corresponding to neuroimaging frames with excessive motion (>1.5mm). Second-level group analyses were conducted by entering the first-level individual models containing the parameter estimates of the four conditions into a mixed-effects analysis, with subject as a random factor.

We then completed region of interest (ROI) analyses by combining data from automated meta-analysis and also a commonly used anatomical brain atlas. This involved combining voxel-wise maps of brain areas involved in reward processing using a mask derived from Neurosynth (Yarkoni et al., 2011), as well as the Harvard-Oxford anatomical atlas available in FSL. Of note, Neurosynth (neurosynth.org), is an automated brain-mapping application that uses text-mining, meta-analysis, and machine-learning techniques to generate a large database of mappings between neural and behavioral/cognitive states. Here, we focused on the term “*value*” from Neurosynth’s past studies database, which included 470 studies. The Harvard-Oxford Cortical and Subcortical Structural Atlases is a probabilistic atlas provided by the Harvard Center for Morphometric Analysis. Only clusters of ≥ 50 voxels were included and explored in proceeding steps. By combining these data sources, we aimed to isolate brain areas related to reward that were anatomically distinct (and did not span multiple areas; derived ROIs shown in *Figure 1*). Such an approach may overcome issues with voxel-wise testing (e.g., clusters of interest spanning multiple discrete brain regions), as well as challenges with exact neural localization (e.g., portions of discrete brain regions not being involved with a candidate neurobehavioral process, such as value-based decision-making) (Woo et al., 2014). We extracted the mean activity for these parametric modulators (i.e., EV and PE) for each ROI for “*approached*” stimuli. Approached stimuli were our primary focus given past reports showing effects of stress on positive feedback and rewards (Hanson et al., 2018, 2016, 2015a). Additional, exploratory analyses related to “avoided” stimuli are detailed in our Supplemental Materials.

**Figure 1.**
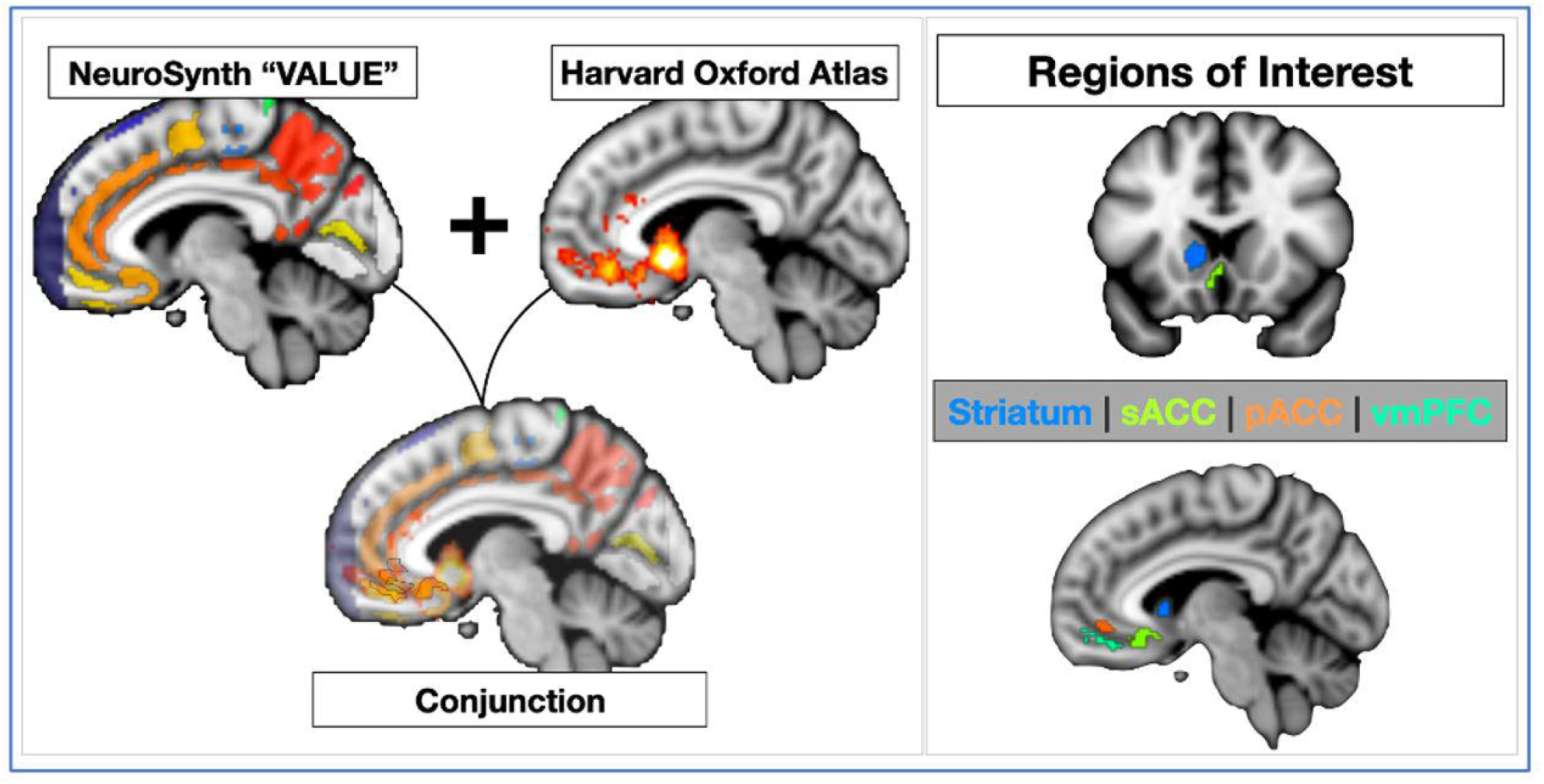
Graphical depiction of how our brain regions of interest were derived. These procedures involved combining automated meta-analyses from NeuroSynth.org related to the search term “value” (shown on the top left) and a commonly used anatomical atlas, the Harvard-Oxford Atlas (shown on the top-right). Combining these sources, we are able to overcome issues with each approach. As shown in the bottom of the figure, this yielded four regions that were ≥50 voxels. These were used in subsequent data reduction steps and analyses.

### Statistical Analysis

Statistical analyses involved examination of neuroimaging measures derived from our passive avoidance learning task. After ROI extraction, linear regression models were constructed using the R statistical package (http://cran.r-project.org). These models examined associations between family income, task-based functional activity for “approached” stimuli, and depressive symptoms. Sex (binary-coded) was included as a covariate in all analyses. First, we examined whether family income (measured when youth approximately 10 and 11 years of age, entered as the independent variable) was related to activity in brain regions derived using the methods described above (entered as the dependent variable). To reduce Type I error, we also adjusted our p-values based on the Benjamini & Hochberg False Discovery Rate Correction (Benjamini and Hochberg, 1995). Next, we tested associations between any brain regions related to childhood family income (as the independent variable, entered in separate models) and adolescent depressive symptoms (measured when youth were approximately 16 years of age, as a dependent variable). We then tested whether family income (X) was associated with later depressive symptoms (Y) and whether the observed association was mediated by individual differences in task-based functional activity (M). This test was done by computing the product of the indirect effects (a X b), as well as the total effect (c + a × b) using bootstrap confidence intervals (95% CIs) based on 5,000 draws with replacement in R’s *lavaan* package; effects were deemed significant if the confidence interval for indirect effects did not include zero (Preacher and Hayes, 2008). Finally, related to our exploratory aims, we tested if associations existed between environmental harshness and unpredictability, and brain activity during this decision-making task using linear regression models (brain activity entered as the dependent variable; harshness or unpredictability entered as an independent variable in separate models).

## 3. Results

### 3.1 Family Income and Neural Markers of Decision-Making Processes

Across our derived sets of ROIs, we investigated connections between prediction error and expected value for portions of the PFC and the striatum. Examining portions of the PFC and EV, multiple anatomically distinct regions related to reward were associated with family income. Specifically, regression analyses indicated that subgenual, as well as more ventromedial, divisions were positively related to this SES measure (sACC *β=0.331, p=0.002*, ventromedial-PFC *β=0.244, p=0.0319*). Put another way, with greater income, greater activity was seen in these regions for stimuli that participants “approached.” Of note, EV-related activity in the pACC for approach stimuli was not related to family income (*β=0.187*, p=.11). With PE and portions of the PFC and EV, regression analysis indicated that activity in the pACC were related to family income (*β=0.267, p=0.021*). Associations for PE in our other PFC ROIs were non-significant (all p’s>.06). In regard to the striatum, family income was not related to brain activity for PE or EV (all p’s>.33). As noted above, given the number of statistical tests conducted (4 ROIs x 2 decision-processes [PE and EV]) and to further reproducibility, we adjusted our p-values based on the Benjamini & Hochberg False Discovery Rate Correction. After application of this correction, only associations between sACC and family income remained significant (*p=0.023; as shown in Figure 2*); all other corrected p-values were ≥.08.

**Figure 2.**
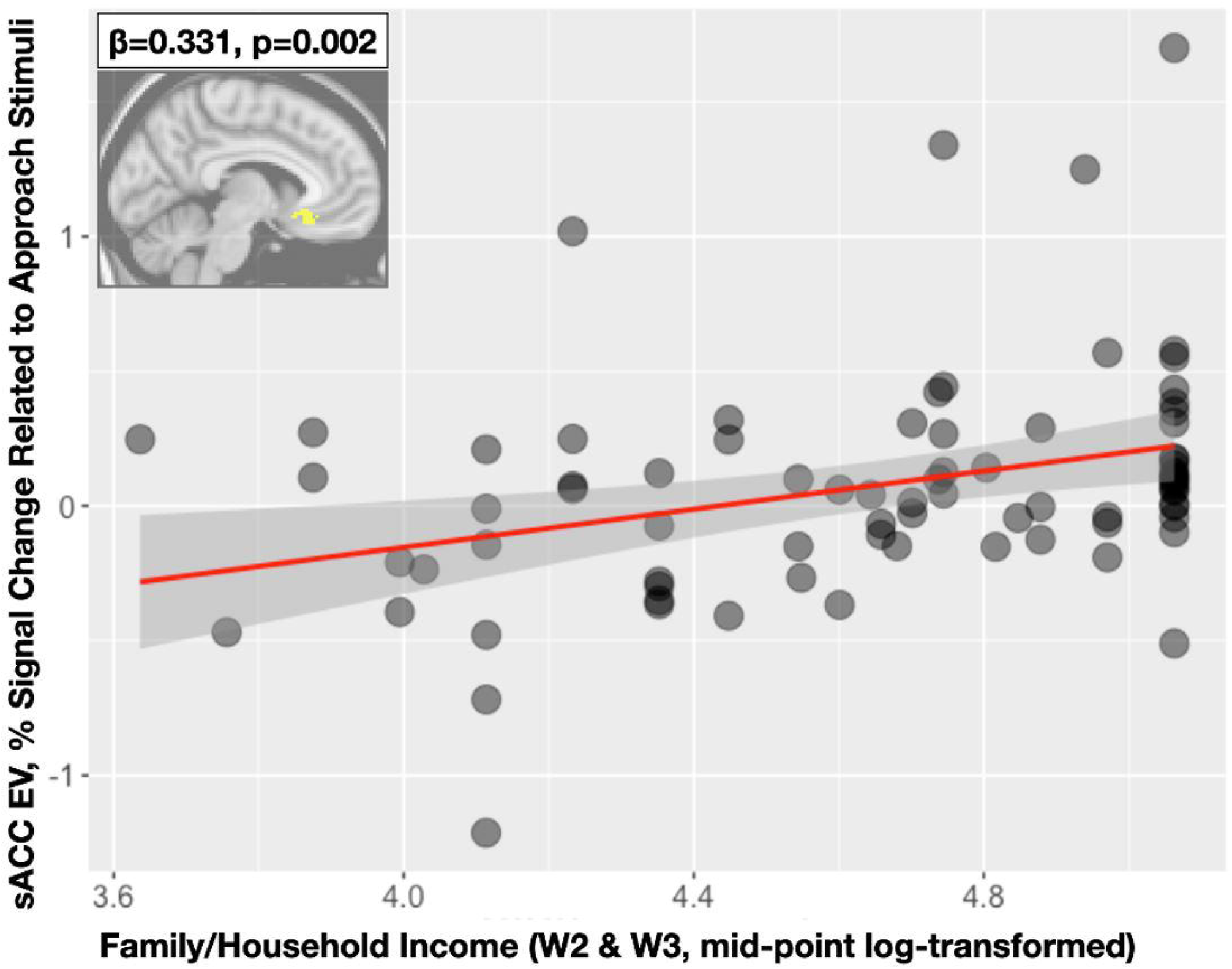
Scatterplot showing associations between early family income (Wave 2 and 3, mid-point log-transformed) shown on the horizontal axis, and fMRI BOLD percent signal change for the subgenual anterior cingulate (sACC) for the trial-wise parametric modulator of expected value on the vertical axis. With greater levels of family income, greater activity is seen in this sACC ROI.

### 3.2 Neural Markers and Depression

We next investigated the potential psychological relevance of these neurobiological variations by examining associations between functional brain activity related to aspects of decision-making and a commonly used, symptom-based measure of depression. In line with our predictions and past work showing corticostriatal differences in depressed samples, we found significant associations between activity in the sACC (for approach EV) and adolescent depression; specifically, higher withdrawn symptoms, approximately one year after the fMRI scan, were correlated with lower sACC activity (*β=-0.269, p=0.017*; as shown in *Figure 3*).

**Figure 3.**
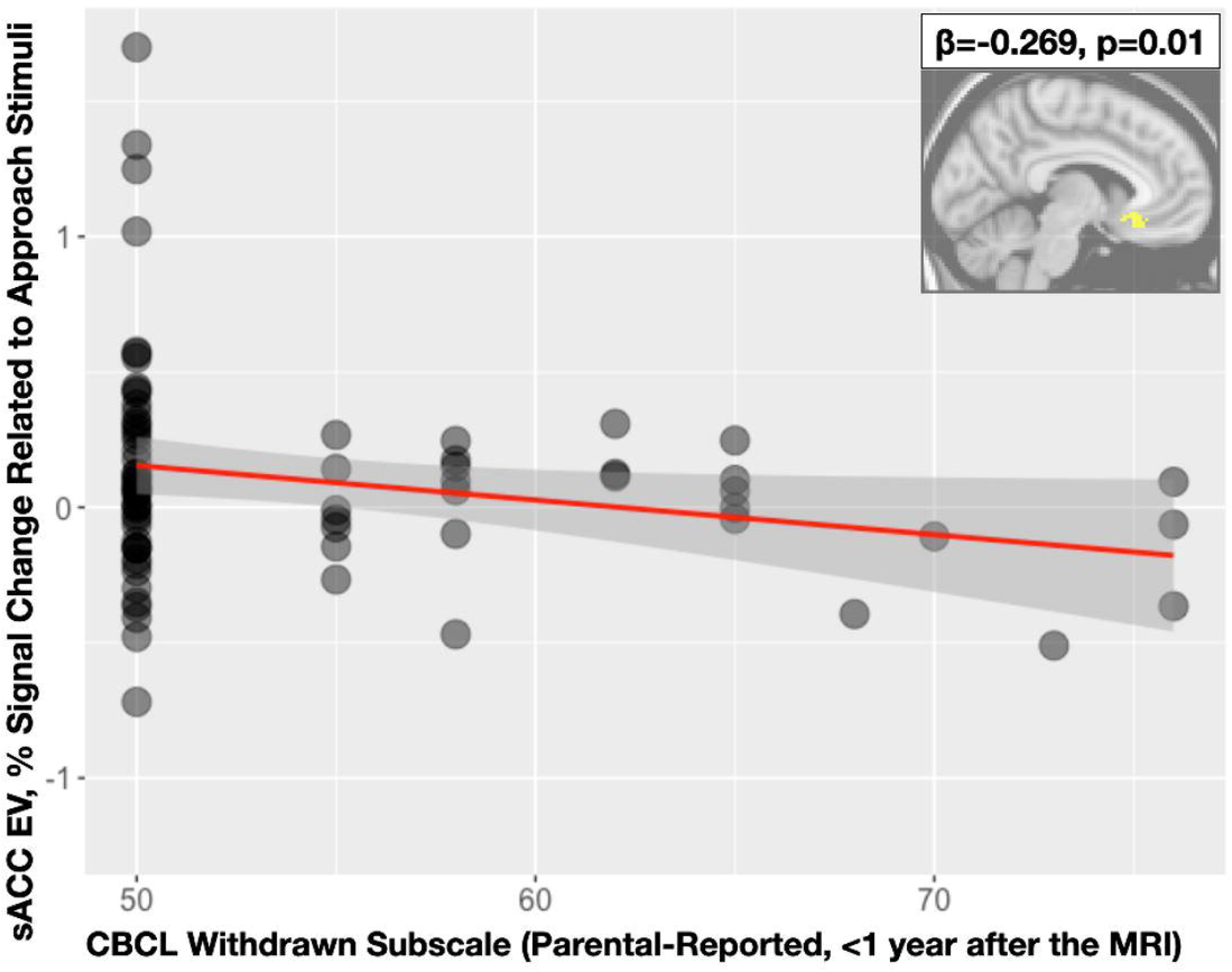
Scatterplot showing associations between parental report of withdrawn symptomatology shown on the horizontal axis, and fMRI BOLD percent signal change for the subgenual anterior cingulate (sACC) for the trial-wise parametric modulator of expected value on the vertical axis. Lower activity in this sACC ROI is related to greater withdrawn symptoms (a proxy for youth depression).

**Figure 4.**
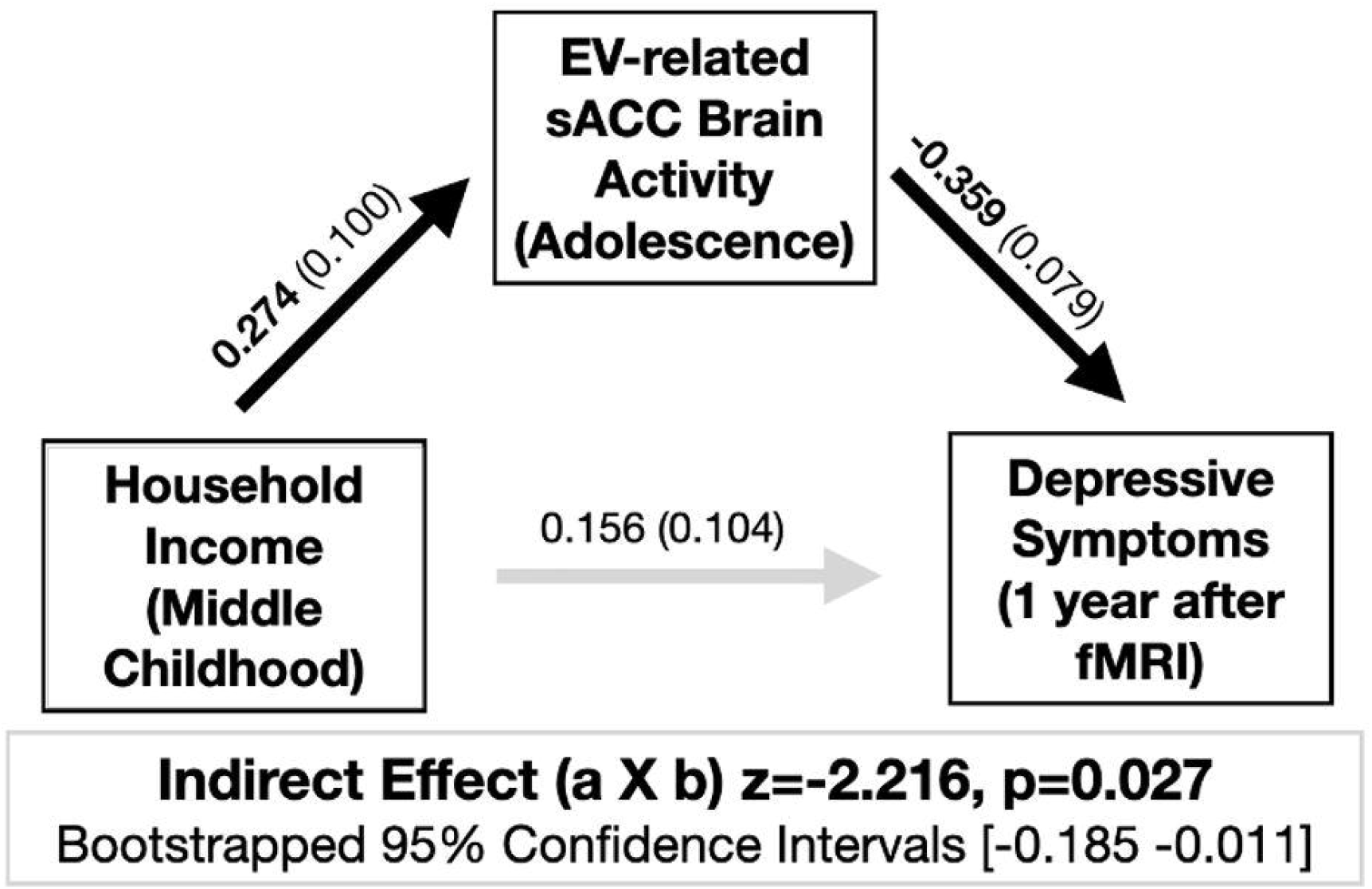
Statistical mediation models are shown in this figure. Standardized regression coefficients along with their standard errors *(in parentheses)* are shown for each path. Significant coefficients are balded (Path a: Income to Brain Activity; Path b: Brain Activity to Depression). The indirect effect [aXb] is significant (p=.027); however, and of important note, the direct effect from income to depression was not (p=0.134).

### 3.3 Probing the Mediating Role of the sACC

Given connections between sACC activity, family income, and depressive symptoms, we tested for potential statistical mediation by entering family income (X), depressive symptoms (Y), and sACC activity, (M) into nonparametric bootstrapped models. Mirroring the results reported above, we find connections between sACC activity and family income (z=2.737, p=0.006), as well as sACC activity and later depressive symptoms (z=-4.536, p<.0001). The indirect effect (combining these two effects, a × b) was significant in the model containing the direct path from family income to depressive symptoms (B=-0.098, SE=0.044, z=-2.216, p= 0.027; 95% CI=-0.185 to - 0.011). However, it should be noted that the direct association between family income and later depressive symptoms was non-significant (p=0.134).

### 3.4 Exploratory Analysis Regarding Dimensions of Adversity

Motivated by past reports from our group (Chang et al., 2019) and connected to different adverse experiences common in impoverished environments, we also explored associations between functional activity and two dimensions of adversity (harshness and unpredictability). Across our ROIs (the subgenual cingulate [sACC], perigenual cingulate [pACC], ventromedial PFC [vmPFC]; striatum) and multiple neural markers (PE, EV), there were no associations with unpredictability or harshness (*all p’s >*. *19, uncorrected*). We also completed Bayesian versions of these models and found similar patterns (as noted in our Supplemental Materials).

## 4. Discussion

Bridging together perspectives from cognitive, developmental, and clinical psychology, as well as neuroscience, the current study examined whether childhood family income was associated with neurobiological functioning related to reward learning and decision-making processes, and whether these alterations were associated with adolescent depression. At the neural level, we found that adolescents with lower childhood family income demonstrated reduced representation of EV in the sACC when making decisions during a learning task. This blunted sACC reactivity was significantly related to greater depressive symptoms one year later. Interestingly, indirect effect/mediation analyses suggested that depressive symptoms were predicted by considering income-related differences in sACC reactivity and also relations between sACC reactivity and depression. Of important note, these effects appear to only be for “approached” stimuli (see our *Supplemental Materials* for non-significant, exploratory analyses for avoided stimuli). Our results suggest unique impacts of rewarded or positive stimuli learning. As such, these findings offer important insights into the effect of lower family income on corticostriatal reactivity, and how these neurobiological changes may in turn give rise to depressive symptoms in adolescence.

Our results showing an association between lower childhood family income and lower sACC activity complement previous findings on early stress exposure and neural alterations within the corticostriatal circuit. For instance, studies consistently find that exposure to child maltreatment, such as physical abuse or social neglect, is associated with reduced reward-related activation in the striatum (Dillon et al., 2009; Hanson et al., 2016, 2015a; Mehta et al., 2010). In terms of the broader corticostriatal circuit, parental education has been related to corticostriatal functionality, with lower education relating to lower activity in portions of the anterior cingulate for rewarded compared to non-rewarded stimuli (Gianaros et al., 2011). These collective findings relate to recent work by Gerin and colleagues that found hypoactivation in portions of the anterior cingulate and vmPFC in maltreated youth compared to their non-maltreated counterparts (Gerin et al., 2017). These neural differences may connect to behavioral impairments in decision-making, such as being insensitive to changes in expected value or speeding responses when the chance of winning increases (Guyer et al., 2006; Weller and Fisher, 2013). Considered more broadly, different adverse exposures and experiences may influence neural and behavioral functioning involved in reward learning and decision-making.

Given our results, we believe it is important to consider how poverty may be impacting neural and behavioral functioning. As detailed in our introduction, poverty may be conceptualized as an adverse exposure, or developmental context, capturing a heightened probability of many deleterious events likely to influence development.

Adverse experiences, in contrast, are the direct measurement of what actually happens to a child (McLaughlin et al., 2020). Connected to these ideas, we attempted to probe different dimensions of experience, harshness versus unpredictability, common to poverty to more deeply understand our results. Experiences of harshness are external threats, while unpredictability involves uncertain and unexpected variations in developmental experiences, particularly unpredictable, chaotic, and harsh events. We, however, did not find any links with brain activity in our ROIs and these dimensions of adversity. Our project had reasonable measures of these constructs, but this was not the original focus of the work. Future research would benefit from the inclusion of these dimensions of adversity to compliment traditional measures of poverty.

Related to dimensions of experience, there are multiple connected, but distinct, conceptual models about unpredictability. Ellis and colleagues (e.g., (Ellis et al., 2009) anchor investigations in an evolutionary frame and define unpredictability as variability in harshness. Baram (e.g., (Baram et al., 2012), using more neurodevelopmental conceptualizations, considers unpredictability more generally, in terms of volatile environmental input and stimuli. We believe this later concept may be particularly important to consider in the “lived experiences” of poverty and the stressors associated with lower family income. Environmental instability may be a powerful factor influencing youth’s well-being, especially during adolescence when the brain is rapidly developing and youth may be navigating new and changing social environments (Blakemore and Mills, 2014; Fuhrmann et al., 2015). This may come in the form of day-to-day unpredictable social interactions (e.g., parental inconsistency)(Davis et al., 2017), as well as unexpected macro-changes in the environment (e.g., residential instability (Adam, 2004). It will be critical for future work in lower socioeconomic status samples to include measures aimed at capturing specific experiences related to poverty. Such targeted investigations, inspired by our findings, may be able to more robustly decompose if brain activity related to reward learning and decision-making connects with different dimensions of adverse childhood experiences.

Reflecting on the significant association between sACC activity and later symptoms of depression, it is important to unpack and think about recent reports focused on this form of psychopathology independent of adversity. For example, heightened connectivity between limbic regions (like the amygdala) and sACC have commonly been reported in samples of individuals with depression, with many arguing this represents a neural risk marker for the development of depression (Marusak et al., 2016). While our results may seem complex (or even potentially counterintuitive) in light of these patterns, it is important to consider: 1) the task domain (or lack thereof) being probed, and 2) challenges with specific spatial locations in the prefrontal cortex. Connectivity-based resting state studies have constituted much of the work finding heightened sACC “activity” (Connolly et al., 2013; Greicius et al., 2007). It is likely that brain activity will show different patterns when using different tasks (e.g., emotion processing versus reward/decision-making). Finally, many studies examine the PFC (or adjacent anterior cingulate cortex), but reported findings are often variable in naming convention and cover large, heterogeneous regions of the brain (Marusak et al., 2016).

Connecting our significant results to previous cognitive neuroscience work, portions of the PFC, including the sACC, are implicated in tracking the expected value of rewards, and these areas of the PFC allow for flexible, adaptive decision making in response to changing stimulus-outcome contingencies (Frank and Claus, 2006; Haber and Knutson, 2010; Murray et al., 2007). Given these findings, we believe it is important to expand neurobiological foci in the search for brain-based mechanisms linking adversity to psychopathology (Palacios-Barrios and Hanson, 2019). The majority of investigations (Hanson et al., 2019, 2015c, 2015b) have focused on the amygdala and the broader corticolimbic circuit. Fewer studies have centered in on corticostriatal circuitry in relation to child poverty and other early stressors (cf. (Romens et al., 2015). With deep relations to multiple forms and aspects of psychopathology (Snyder et al., 2017), in-depth investigations of corticostriatal networks could aid in understanding the impact of negative environmental experiences. Taking cues from affective neuroscience, many research groups have gained additional purchase in understanding depression (independent of adversity) by focusing on aspects of reward learning and decision-making. For example, different research groups have examined reversal learning where stimulus-reward pairings are learned and then contingencies are switched (Cools et al., 2002; Fellows and Farah, 2003). Behavioral and neurobiological differences have been noted between individuals with and without depression (Remijnse et al., 2009; Robinson et al., 2012), and this could be a particularly fruitful avenue of investigation in relation to adversity-related psychopathology.

Considering the strengths of our work, the current study benefitted from the use of a prospective, longitudinal study design that permitted temporally separated assessment of childhood family income, adolescent brain activity, and later depressive symptoms. Recruitment and tracking of participants from childhood to adolescence strengthened developmental considerations, as youth during this period experience extensive neurodevelopment of corticostriatal structures, as well as a significantly elevated risk for depression. The reasonable sample size and use of a well-validated reward learning and decision-making task served as additional advantages. However, we must consider potential issues with the work; one notable limitation is the timing of our measurements. Our earliest income variable is when youth were ∼10 years of age, but exposures and experiences in early childhood may profoundly shape development. It is likely that different critical and sensitive periods exists for the impact of poverty, harshness, unpredictability, and other adversities (Gabard-Durnam and McLaughlin, 2020). However, the design of our work is unable to speak to this issue and may represent an overestimation of the effect of income in middle childhood. Future work should aim to have measures of poverty and socioeconomic status earlier in development, while also considering important developmental moderators (e.g., parenting or peer relationships). It will similarly be important to think about what other factors may be driving changes in the brain, such as changes in inflammation and hypothalamic-pituitary-adrenal axis functioning (Kraynak et al., 2019).

These limitations notwithstanding, our results provide suggestive data about potential neurobiological alterations seen after lower family income (and exposure to child poverty). Few investigations have examined the corticostriatal circuit in relation to variables such as family income or other markers of poverty. Even fewer reports have leveraged novel paradigms and approaches derived from basic cognitive neuroscience and neuroeconomics. By decomposing elements of reward learning and decision-making, we uncovered novel connections with child poverty, brain activity, and adolescent depression. These neural alterations may be a potential mechanism underlying the commonly seen associations between child poverty and later depression. Variations within portions of PFC may convey risk for depression and other facets of mood dysregulation (Lemogne et al., 2012). Indeed, advances in interventions for depression have actually started specifically focusing on cultivating positive affect and reducing anhedonia (Fava and Tomba, 2009; McMakin et al., 2011). Continued progress in this space could have implications for the development and implementation of novel resilience-promoting interventions in those exposed to child poverty and other early life adversities. Similar approaches might be effective in reducing the negative developmental consequences of poverty and the stressors associated with poverty. Additional research is needed to clarify the complex relations among early poverty exposure and long-term mental health difficulties; our data are, however, a needed step in the ability to predict, prevent, and treat stress-related psychopathology.

