## Supplemental Materials for "Lower neural value signaling in the prefrontal cortex is related to childhood family income and depressive symptomatology during adolescence"

*Additional Information Regarding fMRI Data Acquisition*

Scanning took place on a 3.0 Tesla General Electric scanner at the Duke-UNC Brain Imaging and Analysis Center. Structural and functional images were acquired during the scanning session. A high-resolution T1-weighted anatomical image was acquired with 162 axial slices using a fast-spoiled gradient echo pulse sequence (repetition time = 7.584 ms; echo time = 2.936 ms; field of view = 256 mm; image matrix = 256×256; voxel size = 1×1×1 mm; flip angle = 121 degrees). The anatomical image was used for normalization and co-registration with the functional data.

*Sensitivity Analyses included fMRI Signal-to-Noise Ratio*

Given challenges in imaging portions of the prefrontal cortex, especially more ventral regions, we ran additional statistical models controlling for signal-to-noise ratio in our regions of interest. This was completed in AFNI using their 3dTstat program and the '-tsnr' flag (for computation of temporal signal to noise ratio). Related to our findings of decreased brain activity for EV in sACC in relation to income, we find that this relation strengthens when signal-to-noise ratio in the sACC is included (*Original Analysis* $\beta=0.331$, $p=0.002$; *With Signal-to-Noise* $\beta=0.376$, $p<0.001$). Associations between EV-related sACC brain activity and withdrawn symptoms were similar in magnitude when accounting for signal-to-noise ratio in the sACC (*Original Analysis* $\beta=-0.269$, $p=0.017$; *With Signal-to-Noise* $\beta= -0.238$, $p=0.03$). Of note, fieldmap scans were not collected in this sample, so formal distortion correction was not employed. However, and related to our questions of interest, the correlation between signal-to-noise ratio in the sACC and EV-related brain activity in the sACC was modest (r=-0.159, p=0.163). As shown in Supplemental Figure 2, signal-to-noise ratio extracted from each of our ROIs was not strongly related to EV-related brain activity in those same brain regions.

*Neural Responsivity (Prediction Error & Expected Value) During "Avoided" Stimuli*

While we were specifically interested in brain activity related to positive (or approached) stimuli, we thought it important to also report patterns related to avoided stimuli. This was to probe whether the regional differences that were significant in the main manuscript were specific to positive/reward stimuli, or whether brain differences might be more general (or non-specific) neural variations. To these ends, we extracted mean activity for parametric modulators (Expected Value and Prediction Error) for "avoided stimuli". This was done for any regions that were associated with income in our main manuscript (with approach stimuli; for EV: sACC and vmPFC; for PE: pACC). We then examined whether family income (measured when youth were approximately 10 and 11 years of age, entered as the independent variable) was related to activity in these brain regions. Interestingly and speaking to potential specificity of the results detailed in the main manuscript, similar associations were not seen between family income and brain activity for avoidance expected value in our ROIs (sACC $\beta=0.024$, $p=0.835$, ventromedial PFC $\beta=-0.045$, $p=0.697$). For the sACC, this difference between correlations was significantly different ($z=1.95$, $p=.05$), suggesting that associations between sACC and family income were unique to approach expected value. This difference between approach- and avoidance-based correlations did not reach significance for the ventromedial-PFC (*z=1.77, p=0.08*). For prediction error, income was not related to brain activity in the pACC ROIs (*β=0.110, p=0.342*). This difference between approach- and avoidance-based correlations, however, did not reach significance for the pACC (*z=1, p=0.32).*

*Examining Connections Between Brain Activity, Family Income, and Adverse Experiences*

As an alternative to a frequentist approach and a singular focus on null hypothesis testing, we also constructed statistical models using a Bayesian Framework (as implemented in the '*bayestestR'* library in R (Makowski et al., 2019). This would allow us, based on the observed data and a prior belief about the result, to calculate a probability distribution ("posterior") of the effect that is compatible with the observed data. In keeping with current best practices, we described the posterior distribution using the median estimate of the effect, as well as the 89% Credible (*confidence*) Interval of the association (Kruschke, 2014). Of note, researchers have suggested that the commonly-used 95% confidence interval might not be appropriate for Bayesian posterior distributions, due to low stability if not enough posterior samples are drawn (Kruschke, 2014).

Paralleling the methods in the main manuscript, we examined if brain activity in regions of interest that were significant for income (with approach stimuli; for EV: sACC and vmPFC; for PE: pACC) were related to different dimensional experiences of adversity (e.g., harshness; unpredictability). These separate models had brain-activity entered as the dependent variable and sex and income entered as independent variables.

For approach EV-related brain activity, neither harshness, nor unpredictability, was related to brain activity in the sACC (harshness Median=0.020, 89% CI= -0.077 to 0.114; unpredictability Median= -0.001, 89% CI=-0.022 to 0.023) or in vmPFC (harshness Median= 0.025, 89% CI=-0.079 to 0.131; unpredictability Median=-0.008, 89% CI=-0.033 to 0.015). For approach PE-related brain activity in the pACC, our Bayesian models suggested no associations between this neural responsivity and dimensions of adversity (harshness Median=0.017, 89% CI=-0.044 to 0.078; unpredictability Median=0.001, 89% CI=-0.013 to 0.015). Of note, Bayesian models did suggest similar connections between family income and brain activity as reported in the main manuscript. Family income was related to approach EV-related brain activity in the sACC (Median=0.355, 89% CI=0.159 to 0.527) and vmPFC (Median=0.295, 89% CI=-0.086 to 0.525), and approach EV-related brain activity in the pACC (Median=0.183, 89% CI=0.051 to 0.302).

*Behavioral Responsivity During the fMRI Learning Task*

To examine behavioral performance on the task, we assessed each participant's omission errors (the proportion of trials in which reward stimuli were avoided), commission errors (the proportion of trials in which punishment stimuli were approached), and total errors (sum of omission and commission errors). This occurred through separate regression model tested whether childhood family income (entered as the independent variable) was associated with omission and commission errors (entered in separate models, as dependent variables).

Regarding behavioral measures derived from the passive-avoidance task, family income was not related to overall reaction time, reaction time for approach or avoidant stimuli, number of overall responses, number of responses for approach or avoidant stimuli, omission errors, or commission errors (*all p's > .1*). Early family income, however, did significantly predict responses to high magnitude rewards ($\beta=-0.269$, $p=0.013$). This result suggests that adolescents with lower childhood family income were significantly less likely to respond to stimuli predicting higher magnitude rewards. Income was not related to responses for low magnitude rewards, low magnitude punishments, or high magnitude punishments (*all p's > .63*).

***Supplemental Figure 1***.

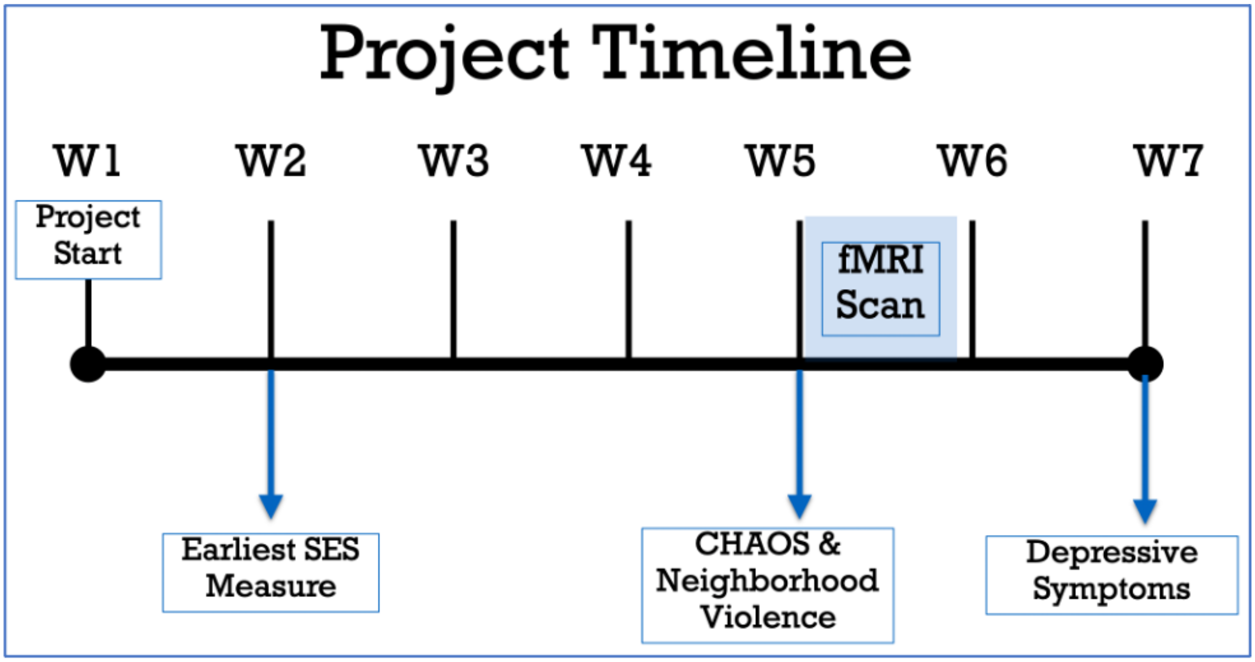


***Figure Caption***: A graphic timeline showing the timing of different study elements, including measures of socioeconomic status, dimensions of adversity, and depressive symptoms. The timeline also shows when fMRI scanning occurred (as shown in the blue shaded box).

***Supplemental Figure 2.***

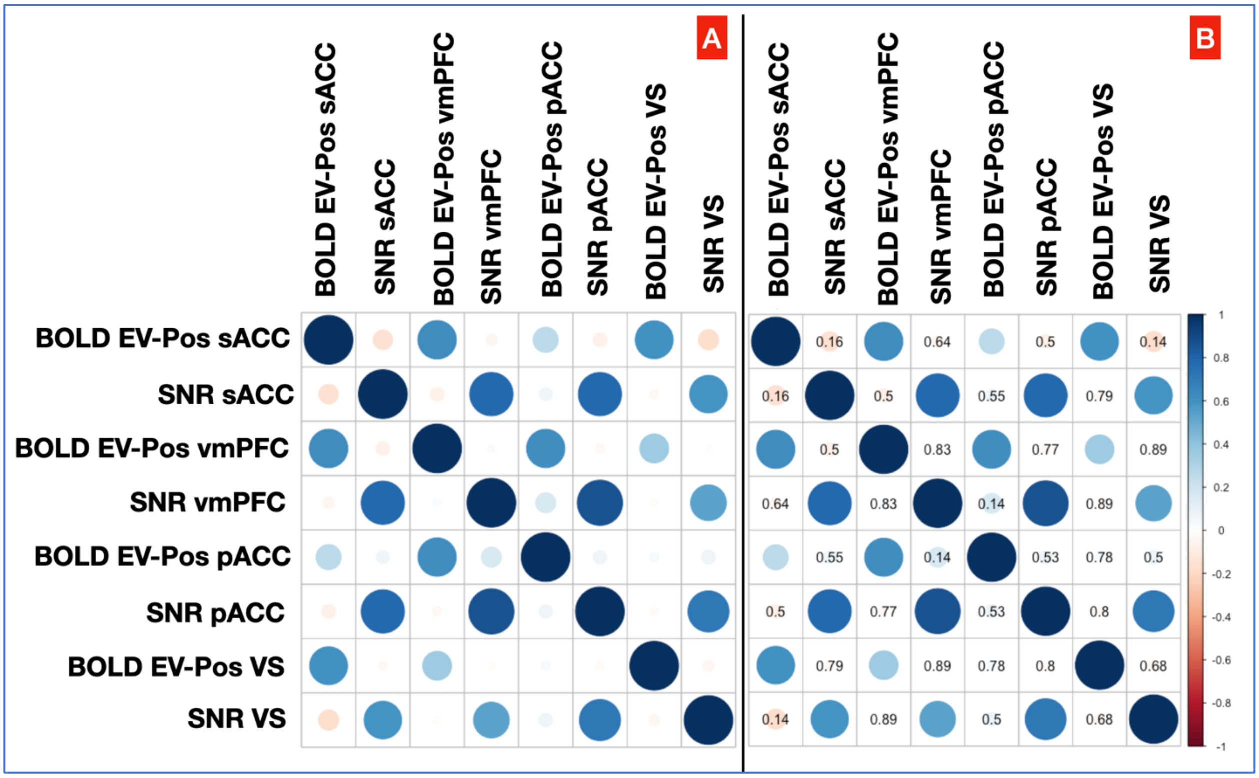


***Figure Caption***: This two panel figure shows correlograms of brain activity for different regions of interest (with the prefix "BOLD") and signal-to-noise ratio (with the prefix "SNR"). The correlograms are identical, the right side (*panel B*) simply showing the p-values of the correlations between brain activity and signal-to-noise ratios (*all non-significant*). All brain activity contrasts were significantly correlated, varying in magnitude from r=.2-.6
